# Detecting and typing *Chlamydia trachomatis* strains in metagenomes using the MetaChlam pipeline

**DOI:** 10.64898/2026.08.18.745514

**Authors:** Parul Sharma, Deborah Dean, Timothy D Read

## Abstract

The Gram negative bacteria *Chlamydia trachomatis* (*Ct*), an obligate intracellular human pathogen, is a predominant cause of sexually transmitted infections and ocular trachoma globally, exerting a significant impact on public health. *Ct* “strains” (major lineages within the species) are known to have different tissue tropisms and be associated with different disease outcomes. Metagenome samples from typical sites where *Ct* infects (e.g., endocervix, conjunctiva, rectum) rarely contain enough reads for traditional genotyping methods such as Multi-Locus Sequence Typing (MLST) or *ompA* genotyping. To overcome these limitations, we implemented an ensemble tool called MetaChlam that can accurately classify *Ct* strains with as few as 250 *Ct* reads. Using 109 publicly available *Ct* genomes from naturally circulating strains, we established that an ANI-based threshold of 99.75% was capable of distinguishing *Ct* strains from each other. We implemented metagenome-based typing using the previously developed LINtax, Strainscan, StrainGE, and Sourmash softwares. MetaChlam integrated the four tools along with custom databases into an automated nextflow pipeline. Using simulated metagenomic reads, we found that our pipeline accurately identified the correct strains in both single strain and multi-strain mixtures of samples. Finally, we showed that MetaChlam had higher specificity for the true presence of *Ct* reads in NCBI SRA metagenomic datasets than NCBI PebbleScout software. A surprising finding of these analyses was that reads from *Ct*, an obligate human intracellular pathogen, can be found as contaminants in samples from sites where the organism is almost certainly not present. Overall, our study enhances the characterization and classification of *Ct* strains and provides protocols for identification and typing of *Ct* in shotgun metagenome data. The MetaChlam pipeline is available on Github: https://github.com/parul-sharma/MetaChlam.

## Introduction

*Chlamydia trachomatis* (*Ct*) is the most common cause of bacterial sexually transmitted infections (STIs), with more than 2.6 million cases in the United States alone (“Sexually Transmitted Infections Surveillance, 2022” 2024). Distinct *Ct* strains cause lymphogranuloma venereum (LGV), which can form ulcers and tends to be a more invasive STI (Haggerty et al. 2010). Other *Ct* strains cause ocular trachoma, the leading cause of infectious blindness worldwide. Given the diverse manifestations of *Ct* infections, understanding the genetic diversity of the organism is crucial for developing effective management and control strategies. Comparative genomic approaches, as illustrated in other species (Viver et al. 2024; Raghuram et al. 2024b), have demonstrated the value of high-resolution strain typing for linking genomic variation to pathogenic potential. Applying similar approaches to *Ct* may clarify how genomic background contributes to its biological and clinical diversity (Joseph et al. 2012; Olagoke et al. 2025; Hadfield et al. 2017).

Several typing techniques have been employed to study the epidemiology and pathogenesis of *Ct* infections. Early efforts focused on serotyping the major outer-membrane protein (MOMP), which was later refined through genotyping the *ompA* gene (Dean et al. 1992). *ompA* genotyping classified *Ct* strains into disease-associated groups: Ocular (A, B, Ba, C), urogenital and anorectal disease (D to K, Da, Ga, Ia, Ja), and LGV (L1 to L3, L2a, L2b, L2c). While *ompA* genotyping is useful in differentiating the major disease lineages, it relies on a single, highly recombinogenic locus. Multilocus sequence typing (MLST) (Jolley et al. 2018; Dean et al. 2009; Klint et al. 2007; Pannekoek et al. 2008) is another widely used molecular technique for characterizing *Ct* strains and addresses some limitations of *ompA* genotyping by leveraging multiple conserved housekeeping genes, which are less prone to recombination (Maiden 2006). Using MLST, and, later whole genome sequencing (WGS), *Ct* has been organized into four strain groups: LGV, ocular, prevalent urogenital and anorectal (P-UA), and non-prevalent urogenital and anorectal (NP-UA) infections (Dean et al. 2009). In most cases, strains with the same *ompA* type are members of the same strain group, except for recombinants where the genomic backbone is distinct from the *omp*A genotype (Harris et al. 2012; Joseph et al. 2012; Joseph et al. 2011).

The increasing use of metagenomics has transformed infectious disease diagnostics and epidemiology, particularly for pathogens like *Ct* that are difficult to culture (Marakeby et al. 2014; Bommana et al. 2022). Metagenomics potentially enables direct detection of *Ct* in clinical samples, without culture, facilitating identification in complex microbial communities, including asymptomatic infections and cases with low pathogen burden. In addition, metagenomics can help identify co-infections and characterize the broader microbial context of the infection, providing insights into factors influencing *Ct* transmission and pathogenesis. In this study, we assessed the correlation of ANI-based classification with established *Ct* typing schemes, and showed that an ANI threshold of 99.75% effectively distinguishes *Ct* strains with different disease etiologies, proving a fast and accurate genome-based classification technique. To support classification directly from metagenomic data, we evaluated four complementary tools (StrainGE (van Dijk et al. 2022), StrainScan (Liao et al. 2023), LINtax (Sharma 2023) and Sourmash (Titus Brown and Irber 2016)). Performance was assessed using simulated metagenomic datasets of single or mixed strain samples with varying pathogen abundances. These tools were integrated into a Nextflow (Di Tommaso et al. 2017) pipeline, called MetaChlam, that executes all classifiers in parallel for efficient and comprehensive metagenome-based identification of *Ct*. We further applied MetaChlam on publicly available SRA metagenomic datasets, identifying evidence of *Ct* infections in 156 samples across 22 BioProjects.

## Results

### 1. ANI threshold of 99.75% classified *Ct* into four major strain groups

A total of 89 publicly available *Ct* complete genome assemblies (accessed in March 2025) were filtered for genome quality. This dataset was supplemented with 20 high-quality reference genomes from a recent study (Olagoke et al. 2025), resulting in a final set of 109 genomes (Supplementary Table 1). Phylogenetic analysis (Fig 1a), resolved four major clades corresponding to previously defined strain groups: LGV (lymphogranuloma venereum), P-UA (prevalent urogenital and anorectal), NP-UA (non-prevalent urogenital and anorectal) and Ocular (Joseph et al. 2023). These clades comprised 23 LGV, 36 P-UA, 37 NP-UA and 13 Ocular genomes. Across the dataset, 21 distinct *ompA* genotypes were identified. The ocular clade included four genotypes - A, B, Ba, C, while the LGV clade comprised six genotypes - L1, L2, L2a, L2b, L2c, L3. In contrast, the P-UA and NP-UA lineages showed overlapping *ompA* genotypes, with P-UA carrying D, Da, E, F, G and Ja, and NP-UA carrying D, G, H, I, Ia, J, Ja, and K. These results aligned with previous studies of *Ct* genome-based phylogeny(Joseph et al. 2023; Olagoke et al. 2025; Hadfield et al. 2017).

**Fig 1.**
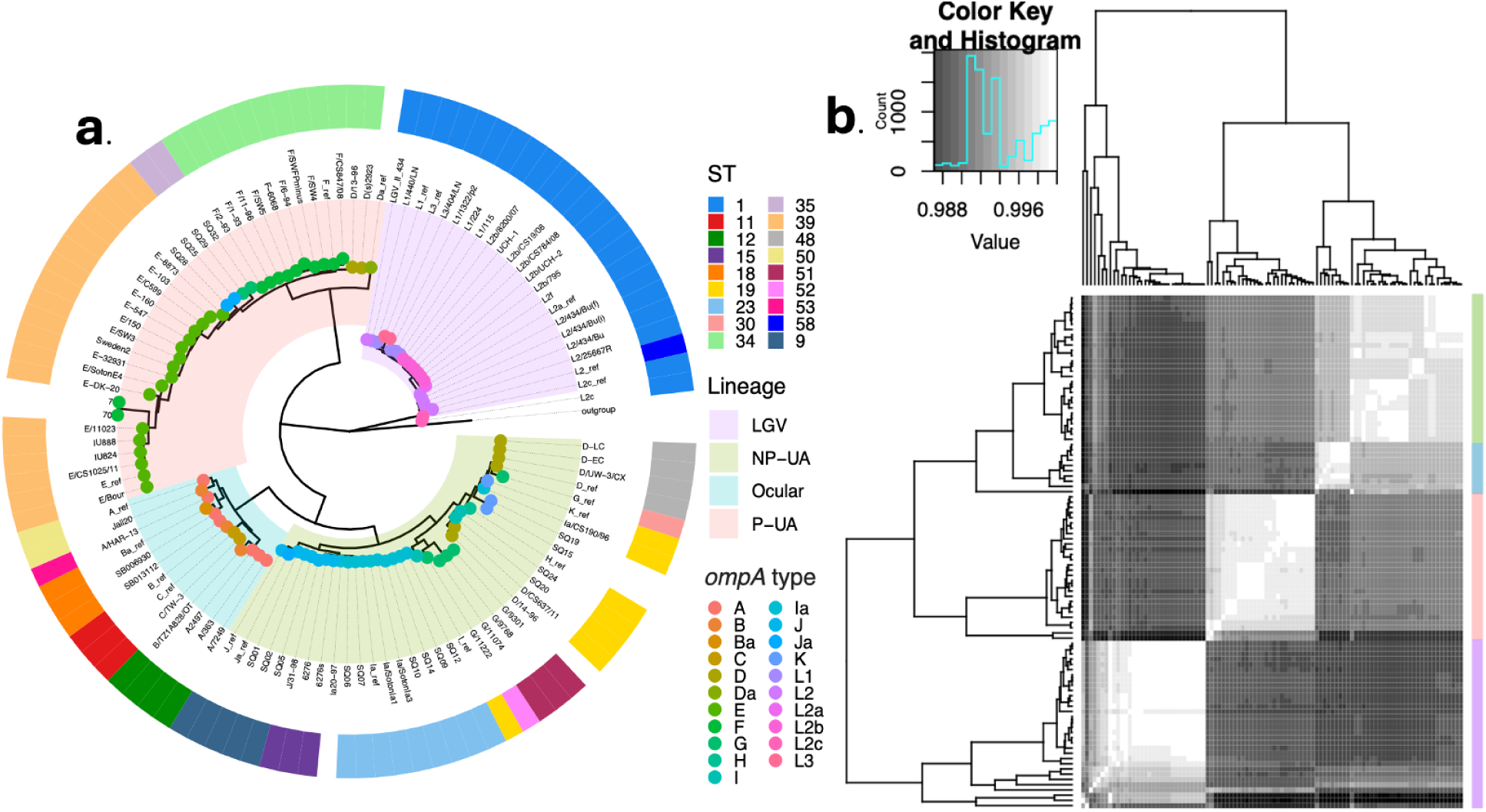
Comparison of trees obtained using core-genome approach and distance based approach. (a) Phylogenetic tree obtained using core-genome analysis of public *Ct* strains from NCBI. All four major lineages are highlighted in different colors representing LGV, NP-UA, P-UA and Ocular strains. Colored dots at the nodes represent the *omp*A genotypes. (b) Heatamp obtained from a distance matrix of all *Ct* strains compared using pairwise-ANI. The Color Key and histogram plot shows the ANI distribution between all pairwise comparisons, and the heatmap highlights the clustering within the strains. Strains grouped together in the same clusters as seen in the core-tree and hence these clusters are marked in the same colors to highlight the 4 lineages.

Average Nucleotide Identity (ANI) (Konstantinidis and Tiedje 2005) analysis compares the entire genomic sequence, providing a more complete view of genetic similarity and divergence, and has been used in numerous comparative genomic studies on other bacteria (Richter and Rosselló-Móra 2009; Jain et al. 2018; Ciufo et al. 2018; Parks et al. 2020). Consistent with the core-genome phylogeny, the ANI-based clustering also partitioned the 109 genomes into the same four clades corresponding to the major infection categories (Fig 1b). Pairwise ANI values ranged from 99.03 to 99.999% (Fig2a). In this paper, we refer to these 4 clades as “strains”, since applying this term to within species groups with the most distinguishing genetic variation provides maximum resolution for “strain-resolved” metagenomic typing (Raghuram et al. 2024). The LGV strain was the most homogenous, with an average pairwise ANI of 99.94 ∓ 0.03%. In contrast, NP-UA, P-UA and Ocular strains showed slightly greater diversity, with average pairwise ANI values of 99.84 ∓ 0.06%, 99.87 ∓ 0.06% and 99.85 ∓ 0.08% respectively (Fig2b).

The Life Identification Number (LIN) (Marakeby et al. 2014; Tian et al. 2020) system builds on ANI to classify and track microbial species at different levels of diversity. Each genome is assigned a unique identifier based on pairwise ANI-thresholds and genomes with similar LINs are grouped into LINgroups, reflecting their genetic relatedness (Vinatzer et al. 2017; Marakeby et al. 2014; Tian et al. 2020). The current LIN implementation clusters genomes based on 20 ANI thresholds ranging from 70-99.999% similarity. Typically, genomes belonging to the same species cluster at a minimum of 95% similarity. The LIN approach includes 13 additional levels of clustering beyond the 95% threshold, which allows for capturing fine-scale genetic differences that traditional taxonomy might overlook (Supplementary Table 1). We found congruence between ANI-based distance trees and maximum likelihood phylogeny based on the core-genome tree (Fig. 2c). We observed that for three strains there were monophyletic clusters at the >99.85% intrastrain ANI that we termed “substrains”. Two stable substrains were identified in the Ocular lineage in both trees. The P-UA and NP-UA strains, exhibiting greater genomic diversity, had four and three substrains respectively. The LGV lineage showed no detectable substrain structure, which is consistent with its high genomic homogeneity. Incongruence was noted for two individual genomes, SQ18 and SQ19, which the core genome phylogenetic tree classified as NP-UA clade 2, while the ANI-based tree placed them in a clade basal to NP-UA clade 1. Given that core genome phylogenetic trees better reflect evolutionary relationships, we considered them as the ground truth, assigning SQ18 and SQ19 to clade 2.

**Fig 2.**
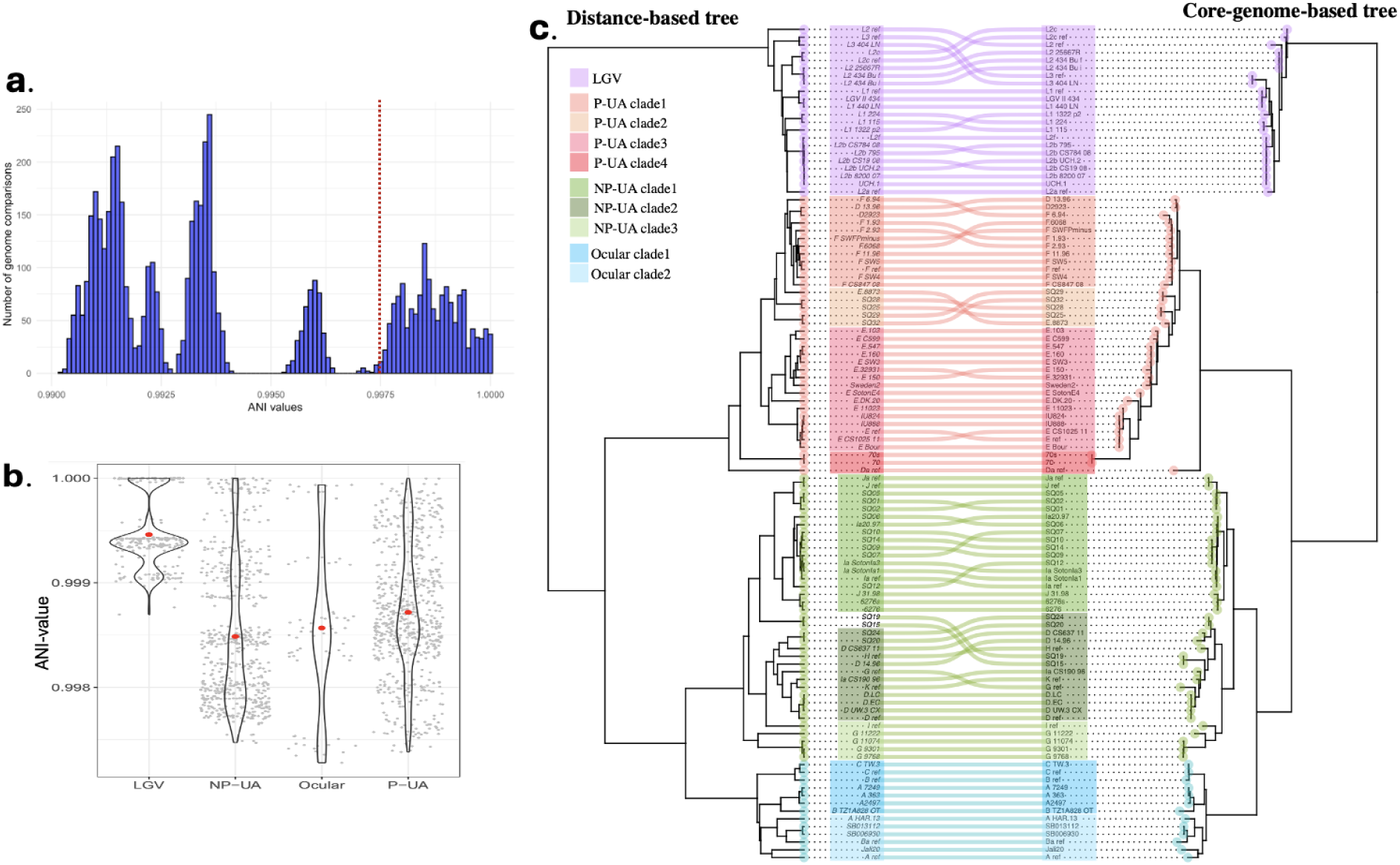
(a) Histogram of pairwise-ANI values among all genome comparisons. The red dotted line represents the strain classification threshold. (b) Pairwise ANI distribution within the LGV, NP-UA, P-UA and Ocular strains. Gray dots represent pairwise ANI values between genomes belonging to the same strain and red dots show the mean. (c) Tanglegram of distance-based tree (on the left) and phylogenetic tree (on the right) shows high congruence between the two approaches. Strains highlighted in the same color scheme between the two trees. Genomes SQ18 and SQ19 of NP-UA strain were not highlighted in the ANI tree (on the left) because of incongruence with the core tree.

### 2. The MetaChlam pipeline: Comparison of tools for identification and classification of *Ct* strains and substrains in metagenome data

We evaluated four previously published tools, StrainGE, Strainscan, Sourmash, and LINtax, for their ability to identify and classify *Ct* genome data to the strain and substrain level. For each tool, reference databases were constructed using the previously described set of 109 genomes. Performance was assessed using simulated Illumina metagenomic datasets. Due to the high genomic similarity within *Ct* strains, with pairwise ANI over 99%, parameters were tuned to ensure high strain-level resolution. For StrainGE, a Jaccard similarity threshold of 99% was required to correctly differentiate the four *Ct* strains. In contrast, StrainScan achieved comparable resolution using its default Jaccard threshold of 95%, highlighting differences in clustering techniques between the tools. Sourmash and LINtax do not rely on explicit clustering thresholds, as their methods inherently accommodate the resolution needed for *Ct* strain differentiation.

We compared the runtime of each tool using a simulated metagenomic sample consisting of one million paired-end reads. Sourmash was the fastest, completing the analysis in just 1.2 seconds, followed by LINtax, with 7.7 seconds. StrainGE and StrainScan required 13.8 seconds and 15.5 seconds, respectively. These results highlight significant differences in computational efficiency, with Sourmash providing the fastest classification for this dataset.

To evaluate the performance of the constituent classifiers incorporated into MetaChlam, we generated simulated metagenomes containing *Ct* at a range of abundances and assessed the ability of each tool to recover the expected strain composition (Supplemental Fig. 1). For single-strain simulations, *Ct* abundance ranged from 20% to 0.005% of the metagenome, corresponding to approximately 100,000 to 25 *Ct* paired-end reads. All four classifiers accurately identified the expected strain at moderate and high abundances, while performance declined as *Ct* abundance decreased. However, the tools differed in their lower limits of detection, with some retaining sensitivity at substantially lower *Ct* abundances than others.

To assess performance in more complex scenarios, we generated simulated metagenomes containing equal mixtures of two *Ct* strains at abundances ranging from 0.5% to 20% of the total metagenome. In these simulations, accuracy was defined as recovery of the complete expected strain combination without missing or spurious strain calls. Mixed-strain simulations proved more challenging than single-strain datasets, particularly at lower *Ct* abundances, and differences between classifiers became more apparent. We further evaluated a complex scenario containing representatives of all four major *Ct* lineages within the same metagenome. Although no single classifier performed optimally across all simulation scenarios, each contributed complementary strengths. These observations motivated the design of MetaChlam as a consensus framework that integrates evidence across multiple classifiers rather than relying on any single method.

All four tools, along with their respective reference databases, were incorporated into the Nextflow-based MetaChlam pipeline. An optional downstream step to infer *ompA* genotypes from classified reads was also included, enabling additional strain-level characterization where coverage permits.

### 3. Thresholds for detection and classification for *Ct* in metagenome data using MetaChlam

Evaluation with the simulated metagenomes (Tables 1-2) revealed that for all non-zero *Ct* abundances, *Ct* detection was highly sensitive, reaching 100% at ≥250 *Ct* reads in single-strain simulations and at all abundances tested in mixed-strain simulations.

**Table 1:** Tiered accuracy of detection with MetaChlam across simulated **single-strain metagenomes** with varying *Ct* read abundances. Each simulated metagenome was composed of 1 million paired-end reads, 125 bp in length. Accuracy values represent the proportion of simulated samples, at each *Ct* read level, for which MetaChlam correctly detected *Ct* and recovered the expected ground-truth. Recovery is shown by support tier based on the number of tools reporting the same correct result (≥1, ≥2, or ≥3 tools). Partial matches and incorrect combinations were counted as incorrect. Value of zero denotes that no classification results were produced at the corresponding read depth.

| Ct reads | N | Ct detected | Strain $\geq 1$ tool | Strain $\geq 2$ tools | Strain $\geq 3$ tools | Substra in $\geq 1$ tool | Substra in $\geq 2$ tools | Substra in $\geq 3$ tools |
| --- | --- | --- | --- | --- | --- | --- | --- | --- |
| Negative Control | 2 | 0 | 0 | 0 | 0 | 0 | 0 | 0 |
| 25 | 113 | 0.18 | 0.18 | 0 | 0 | 0 | 0 | 0 |
| 125 | 113 | 0.47 | 0.47 | 0.01 | 0 | 0.01 | 0 | 0 |
| 250 | 113 | 1 | 1 | 0.58 | 0 | 0.87 | 0 | 0 |
| 500 | 113 | 1 | 1 | 0.96 | 0.64 | 0.98 | 0.76 | 0 |
| 750 | 113 | 1 | 1 | 0.98 | 0.76 | 0.97 | 0.93 | 0.03 |
| 1000 | 113 | 1 | 1 | 1 | 0.77 | 0.98 | 0.95 | 0.01 |
| 1250 | 113 | 1 | 1 | 1 | 0.86 | 0.97 | 0.96 | 0.02 |
| 2500 | 113 | 1 | 1 | 1 | 1 | 0.99 | 0.96 | 0.4 |
| 12500 | 113 | 1 | 1 | 1 | 1 | 0.99 | 0.96 | 0.6 |
| 50000 | 113 | 1 | 1 | 1 | 1 | 0.99 | 0.96 | 0.6 |

**Table 2:** Tiered accuracy of detection with MetaChlam across simulated **mixed-strain metagenomes** with varying *Ct* read abundances. Each simulated metagenome was composed of 1 million paired-end reads, 125 bp in length. Accuracy values represent the proportion of simulated samples, at each *Ct* read level, for which MetaChlam correctly detected *Ct* and recovered the expected ground-truth strain combination or substrain combination. For mixed-strain simulations, a classification was considered correct only if all expected strains were detected; partial matches and incorrect strain combinations were counted as incorrect. Recovery is shown by support tier based on the number of tools reporting the same correct result (≥1, ≥2, or ≥3 tools). Partial matches and incorrect combinations were counted as incorrect. Value of zero denotes that no classification results were produced at the corresponding read depth.

| Ct reads | N | Ct detected | Strain $\geq 1$ tool | Strain $\geq 2$ tools | Strain $\geq 3$ tools | Substrain $\geq 1$ tool | Substrain $\geq 2$ tools | Substrain $\geq 3$ tools |
| --- | --- | --- | --- | --- | --- | --- | --- | --- |
| 0 | 2 | 0 | 0 | 0 | 0 | 0 | 0 | 0 |
| 2500 | 100 | 1 | 0.99 | 0.88 | 0.5 | 0.89 | 0.41 | 0 |
| 5000 | 100 | 1 | 1 | 0.99 | 0.94 | 0.89 | 0.66 | 0.16 |
| 12500 | 100 | 1 | 1 | 1 | 0.99 | 0.97 | 0.83 | 0.01 |
| 25000 | 100 | 1 | 1 | 1 | 1 | 0.99 | 0.92 | 0.41 |
| 50000 | 100 | 1 | 1 | 1 | 1 | 0.99 | 0.91 | 0.42 |

In single-strain metagenomes, strain-level classification improved rapidly with increasing *Ct* read abundance. At 250 *Ct* reads, at least one classifier correctly identified the expected lineage in all simulations, although agreement among classifiers remained limited. Two-tool consensus increased from 58% at 250 *Ct* reads to 96% at 500 *Ct* reads and reached 100% at ≥1,000 *Ct* reads. Three-tool consensus was first observed at 500 *Ct* reads (64%) and increased to 100% at ≥2,500 *Ct* reads. Substrain-level classification required greater read support. At least one classifier correctly identified the expected substrain in 87% of simulations at 250 *Ct* reads and in 97-99% of simulations at ≥500 *Ct* reads. Two-tool substrain consensus reached 76% at 500 *Ct* reads and increased to 95–96% at ≥1,000 *Ct* reads.

Mixed-strain metagenomes were inherently more challenging because successful classification required recovery of the complete expected strain combination. At 2,500 *Ct* reads, at least one classifier correctly recovered the expected strain combination in 99% of simulations, increasing to 100% at ≥5,000 *Ct* reads. Two-tool consensus improved from 88% at 2,500 *Ct* reads to 99–100% at ≥5,000 *Ct* reads, while three-tool consensus reached 94% at 5,000 *Ct* reads and 100% at ≥25,000 *Ct* reads. Substrain-level recovery was more demanding, with two-tool consensus increasing from 41% at 2,500 *Ct* reads to 91-92% at ≥25,000 *Ct* reads.

Based on these results, we proposed a tiered confidence framework for interpreting MetaChlam classifications. *Ct* detection can be considered highly reliable once at least 250 *Ct* reads are recovered. Strain-level assignments supported by two or more classifiers were consistently accurate at ≥1,000 *Ct* reads for single-strain infections and ≥5,000 *Ct* reads for mixed-strain infections. Substrain-level classifications were most reliable when supported by multiple classifiers and sufficient *Ct* read depth, with two-tool consensus exceeding 95% in single-strain simulations at ≥1,000 *Ct* reads and 90% in mixed-strain simulations at ≥25,000 *Ct* reads. These thresholds provide practical guidance for determining the highest level of classification that can be confidently supported in metagenomic datasets.

### 4. *Ct* identification in clinical metagenome samples

The MetaChlam pipeline was evaluated on three publicly available vaginal metagenomic samples from a published study (Table 3), for which the *Ct* genome had also been sequenced using Agilent SureSelect targeted sequencing (Bommana et al. 2022; Joseph et al. 2023). The three samples (98V, 192V and 362V) were obtained from Fijian women who tested positive for *Ct* infection. The number of classified *Ct* reads in these metagenomes represented only 0.01-0.03% of the total reads, corresponding to 513-1533 classified reads.

**Table 3:** Summarization of results for clinical sample

| Data from original study |  |  |  | MetaChlam results |  |  |  |  |
| --- | --- | --- | --- | --- | --- | --- | --- | --- |
| Sample | ID | strain | *ompA* | # *Ct* reads | % *Ct* reads | strain | substrain | *ompA* |
| SRR18765403 | 362V | NP-UA | F | 513 | 0.01 | P-UA/<br>NP-UA | - | - |
| SRR18765393 | 98V | NP-UA | G | 1533 | 0.03 | NP-UA | NP-UA_clade3 | G |
| SRR18765399 | 192V | P-UA | Ja | 1188 | 0.03 | P-UA | P-UA_clade2 /<br>clade1 | - |

MetaChlam produced reliable strain-level classifications, consistent with the tiered detection thresholds. Sample 98V, with 1533 *Ct* classified reads, was consistently identified as belonging to the NP-UA lineage, in agreement with the original study. Additionally, MetaChlam correctly inferred the *omp*A genotype (type G) of the sample and identified the correct clade (NP-UA clade3). For sample 192V, with 1188 *Ct* classified reads, all tools correctly assigned it to the P-UA lineage. Substrain-level classification, however, was inconsistent, with StrainGE and Sourmash identifying P-UA clade2 while StrainScan identifying clade1. As expected based on the proposed thresholds, *ompA* typing could not be reliably supported at this read depth. In contrast, sample 362V contained only 513 *Ct* classified reads, a range below the threshold for reliable strain-level interpretation. Accordingly, MetaChlam produced inconclusive results: LINtax assigned the sample to the P-UA lineage, StrainGE and Sourmash assigned it to the NP-UA lineage, and StrainScan did not report any *Ct* hits. Together, these clinical results validate the tiered detection thresholds derived from simulated data and demonstrate that MetaChlam could recover *Ct* strain, substrain, and also in some cases, *ompA* type, from real metagenomic samples when sufficient *Ct* read depth was available.

### 5. *Ct* identification and typing of *Ct* in public metagenome data

To assess the prevalence and diversity of *Ct* in publicly available data, we screened metagenomic datasets deposited in the NCBI Sequence Read Archive (SRA). Because *Ct* is relatively rare in metagenomic samples, we used the NCBI web service, Pebblescout (Shiryev and Agarwala 2024), to prescreen putative positive samples with varying competition of reads. Pebblescout enables rapid querying of large-scale sequence repositories by indexing k-mers and reporting coverage as the percentage of query k-mers matched in the subject. Using the *Ct* reference strain (D/UW-3/CX, accession:NC_000117.1) as a query (date of query: November 2024), Pebblescout identified ∼66,000 SRA submissions (out of 100,000 queried) with non-zero percentage of reference coverage. The top 500 highest scoring candidate datasets were subsequently selected for downstream analysis with MetaChlam (Supplementary Table 2).

MetaChlam detected *Ct* reads among 158 of the 500 candidate datasets with high confidence classifications (Supplementary Table 3). Among the classified, 119 samples were identified as single-strain *Ct* infections and 39 as mixed-strain infections. For this analysis, a sample was classified as multi-strain if at least two tools independently detected two or more of the same *Ct* strains. An additional 34 datasets contained detectable *Ct* reads but lacked sufficient multi-tool support for confident strain assignment and were therefore classified as low-support detections, while 303 datasets remained unclassified (with less than 250 reads detected but not enough for strain classifications) whereas five datasets contained no detectable *Ct* reads (Table 4).

**Table 4:** MetaChlam results summary on analysis of 500 SRA metagenome entries, categorizing samples into single-strain, multi-strain, and other classifications. *Single-strain infections* are defined as samples where only one *Ct* strain is confidently identified. *Multi-strain infections* indicate the presence of more than one *Ct* strain in the sample. The low-support detection is where only one tool in the pipeline detected a single strain. *These totals contain 96 samples from WGS studies incorrectly submitted as metagenomic datasets.

| MetaChlam classification | Number of Samples |
| --- | --- |
| Single-strain | 119* |
| Mixed-strain | 39* |
| Low-support detection | 34 |
| Unclassified | 303 |
| No *Ct* reads detected | 5 |
| **Total** | **500** |

We found that the number of classified *Ct* reads varied widely, ranging from 221 reads to 42 million reads, with a median read count of 278k per sample. Together, these results demonstrate the robustness of MetaChlam in identifying and characterizing *Ct* infections in complex, real-world metagenomic datasets. These 158 samples originated from 26 distinct Bioprojects (Supplementary Table 4).

The most common *Ct* strain group identified was P-UA, which was predominantly associated with BioProject PRJEB35886, a *Ct* phylogenomic study with 76 samples. Another large study with 20 samples (Bioproject PRJEB34194), studying the infected eye microbiome, exclusively contained the Ocular strains. Both these studies turned out to be *Ct* genome sequencing projects that were mislabelled as metagenome samples. Besides these, 15 samples from Bioproject PRJEB34871 isolated from pooled human fecal samples also contained the P-UA strains. Twenty one samples from seven Bioprojects (PRJNA454826, PRJNA512250, PRJNA608678, PRJNA637785, PRJNA657309, PRJNA730640. PRJNA675301), all isolated from the human or mouse or pig gut were infected with either NP-UA or ocular strains or both (Supplementary Table 4). All human-associated metagenomes with single strain *Ct* detections, were dominated by urogenital (P-UA/NP-UA) lineages (45/45). In contrast, non-human and environmental datasets contained either single-strain ocular or LGV infections (11/17) or mixed infections involving ocular and NP-UA lineages (6/17), resulting in a lineage distribution significantly different from that observed in human-associated metagenomes (Fisher’s exact test, *p* = 2.4 × 10⁻⁸). Metadata of the sample types and study descriptions across *Ct*-positive datasets are visualized as a word cloud in Supplemental Fig 2.

### 6. Ct reads are a contaminant in some metagenomic datasets

Our analysis of public data also uncovered eight samples derived from unconventional sources in which significant numbers of *Ct* reads were detected. These were: (a) four from a project of 45 metagenomic samples from the gut of farmed pigs (PRJNA512250) (He et al. 2021)., contained 2.2k -3k *Ct* reads; (b) three of 18 samples from a deep-sea hydrothermal vent (PRJNA546572) (Reysenbach et al. 2020) with 292, 7.4k and 11.1k reads each; (c) one of eight samples from a saltwater aquarium tank (PRJNA225954) (Kelly et al. 2014) with 1,946 reads. Although unexpected, these findings provided an opportunity to evaluate MetaChlam’s ability to distinguish *Ct* from closely related *Chlamydia* species. Multiple independent validation approaches confirmed that these signals represent *Ct*-derived reads rather than spurious classifications. In the absence of biological context supporting *Ct* in these environments, these detections are best explained as contamination, providing direct evidence that *Ct* sequences can occur as contaminants in metagenomic data.

In considering the samples from project PRJNA512250, we noted pigs are natural hosts of *Chlamydia suis*, a species closely related to *Ct* (average pairwise ANI of ∼79.5%, Supplementary Table 5) and historically considered a biovar of *Ct* (Häcker 2024; Schautteet and Vanrompay 2011). Given this, we hypothesized that MetaChlam may have misclassified *C. suis* as *Ct* in the pig-associated samples. To investigate, we examined metagenome-assembled-genomes (MAGs) reconstructed from these samples. ANI analysis revealed a substantially higher sequence identity with *Ct* than with *C. suis*. The percentage identity between the longest assembled MAG (193 kb in size) from the 4 pig samples and the best *Ct* reference (Ocular *ompA* type A reference strain) was 99.18%, compared to only 80.71% against *C.suis* (Supplementary Table 5). Alignment of the MAG to the *Ct* reference genome showed high sequence similarity and even coverage, with most contigs aligning at ≥95% identity (Fig 4a1-a2). Phylogenetic analysis further placed the MAGs within the Ocular strain cluster of *Ct* (Fig 4a3). The mean depth of coverage across the *Ct* reference genome was 0.65x, corresponding to an expected genome breadth of 47.8% under the Lander-Waterman model (Lander and Waterman 1988). The observed breadth was 45.7%, closely matching theoretical expectations and supporting genome-wide, low-abundance representation rather than localized or artifactual alignment. These results confirm that the detected sequences are derived from *Ct* rather than *C. suis*. However, given the absence of biological context supporting *Ct* in pig gut environments, these signals are most consistent with contamination rather than true host-associated infection or colonization.

**Fig 3:**
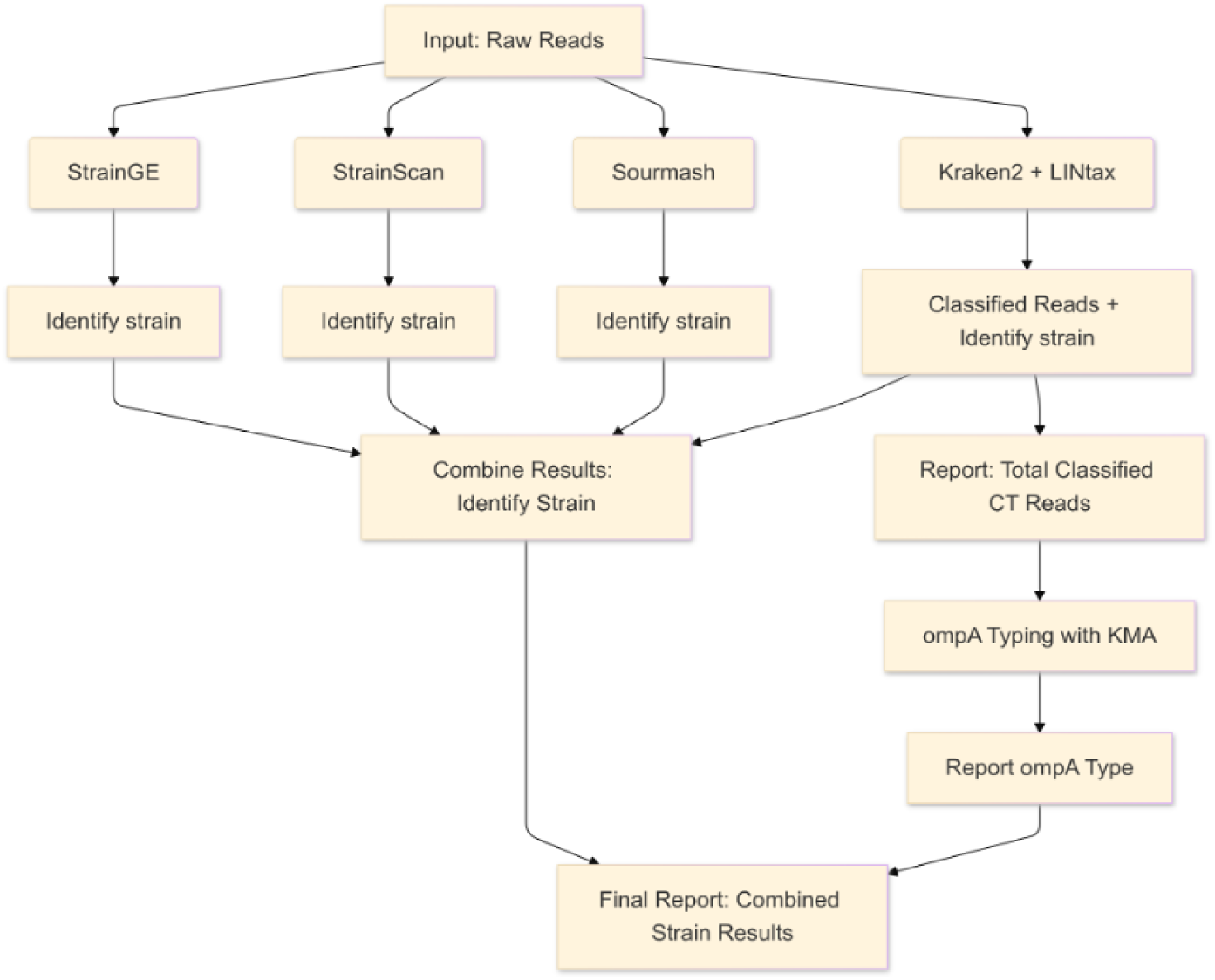
Schematic flow chart for MetaChlam Pipeline. The pipeline is currently tested only on Illumina sequencing reads, which are the most common technology for shotgun metagenomics.

**Fig 4:**
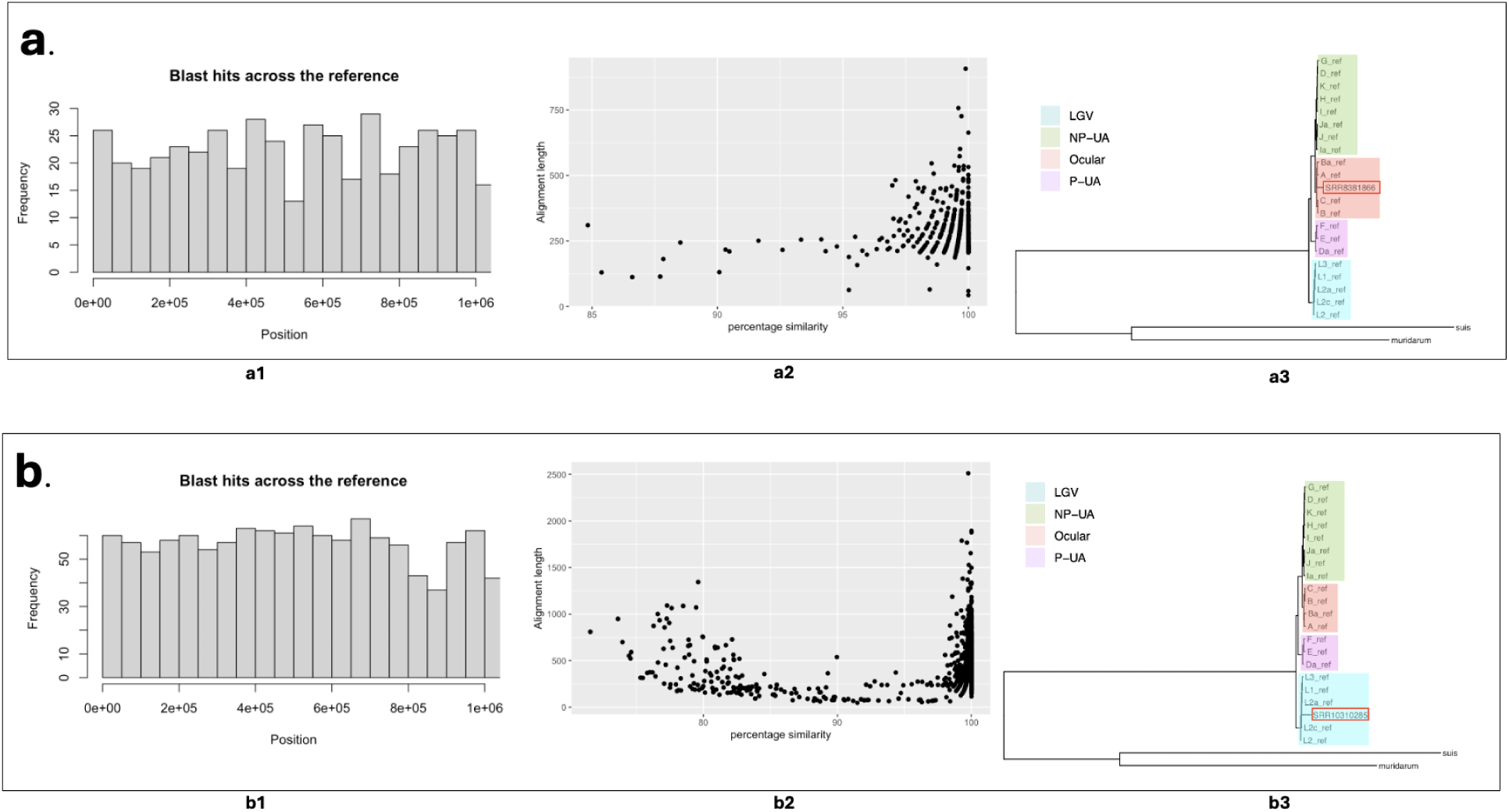
Examples of validation of MetaChlam detection of *Ct* in metagenomes from unexpected sources. (a) Validation using a metagenome-assembled genome (MAG) from pig gut samples: (a1) even genome coverage obtained after aligning *Ct* identified reads from the sample against the reference strain (*omp*A genotype A), (a2) high sequence similarity between assembled contigs and the reference genome, and (a3) phylogenetic analysis with *Ct* reference genomes places the MAG in the ocular strain cluster, confirming detection as an ocular strain. (b) Validation using a MAG from the hydrothermal sample: (b1) even genome coverage obtained after aligning *Ct* identified reads from the sample against the reference strain (ompA genotype L2), (b2) high percentage similarity between assembled contigs and the reference genome, and (b3) phylogenetic analysis with *Ct* reference genomes places the MAG in the LGV strain cluster, confirming detection as a LGV strain.

Detection of *Ct* in metagenomes derived from a saltwater aquarium and deep-sea hydrothermal vents was even more unexpected, as such environments are unlikely to naturally harbor *Ct*. These findings prompted an investigation into potential human-derived contamination. Taxonomic profiling with Kraken2 (Wood et al. 2019) revealed taxa typical of these extreme environments, such as *Sulfurimonas*, *Oceanitermus* and *Persephonella*. Nevertheless, human-derived reads were detected in all 3 samples, constituting 0.28% (690k) and 0.8% (288k) reads in the two hydrothermal vent samples, and 6.6% (82k) reads in the one aquarium sample. MetaChlam identified LGV strain (and ompA type L2) in all three samples. Further examination using MAGs reconstructed from these samples supported MetaChlam’s results. The hydrothermal vent samples yielded relatively complete *Ct* MAGs, with sizes of 698 kb and 554 kb, compared to 198 kb for the aquarium sample. ANI analysis revealed high sequence identity to the *Ct* LGV L2 reference strain, with values of 99.34% and 98.5% for the hydrothermal vent MAGs and 99.86% for the aquarium MAG. Whole-genome alignment demonstrated high sequence similarity, with the majority of contigs aligning at ≥95% identity (Fig. 4b1–b2). The mean depth was 3.17x, for which the Lander-Waterman model of a perfect genome project (Lander and Waterman 1988) predicted 95.8% breadth of coverage. The observed breadth was 75.4%, indicating significant genome-wide representation, although somewhat lower than the theoretical expectation, which could have caused uneven coverage patterns typical of some metagenomic projects. Phylogenetic reconstruction clustered these MAGs with known LGV strains (Fig. 4b3), corroborating MetaChlam’s strain-level classification.

## Discussion

Our analysis established an ANI threshold of 99.75% as effective for differentiating between the four *Ct* strain groups, Ocular, NP-UA, P-UA, and LGV. This threshold is consistent with recent large-scale studies such as a natural ANI gap at 99.2 to 99.8% in within-species analysis of ∼18,000 genomes (Rodriguez-R et al. 2024; Aldeguer-Riquelme et al. 2024), and the use of an 99.5% ANI threshold to differentiate *Staphylococcus aureus* strains (Raghuram et al. 2024a). These consistent results illustrate the broad applicability of ANI thresholds for intraspecies differentiation and reinforce their utility for strain-level genomic analyses. We incorporated the 99.75% ANI threshold into construction of custom reference databases for MetaChlam, enabling consistent strain discrimination across tools. We also introduce a second threshold, at > 99.85% ANI, that we call “substrain” that is useful in distinguishing monophyletic clades within three of the strains (although typing was slightly less accurate than the strain threshold). Justification for the use of the term “strain” follows that outlined in Raghuram et al., 2024(Raghuram et al. 2024). Essentially, to allow the most-effective “strain-resolved” metagenomic typing we advocate for assigning strain to mean the highest level natural sub-species partition.

The MetaChlam pipeline incorporates four complementary tools—LINtax, StrainGE, StrainScan, and Sourmash—each of which uses a different operational definition of a bacterial strain. LINtax groups genomes into epidemiologically meaningful intraspecies clusters, whereas StrainGE, StrainScan, and Sourmash identify the closest matching reference genome based on distinct sequence comparison strategies. Rather than relying on a single classifier, MetaChlam combines evidence across all four tools to generate consensus strain calls. This ensemble strategy leverages the complementary strengths of the individual methods while reducing the influence of tool-specific biases and false-positive classifications. By requiring concordance among multiple classifiers before reporting mixed infections or high-confidence strain assignments, MetaChlam provides a conservative yet sensitive framework for interpreting *Ct* metagenomic data across a broad range of read depths and sample complexities. The simulation study further enabled empirically derived confidence thresholds based on *Ct* read abundance, allowing strain classifications from real metagenomic datasets to be interpreted in the context of expected classification accuracy (Ostvar and Eftekhari Moghadam 2020; Lu et al. 2024).

MetaChlam’s low detection threshold is facilitated by the high genomic conservation of *Ct,* which allows strain discrimination with relatively sparse read support compared to more diverse bacterial taxa (Harris et al. 2012; Joseph et al. 2012). However, read count alone does not determine classification confidence, as samples with similar numbers of *Ct* reads may capture different regions of the genome. Hence, the proposed read thresholds should be viewed as empirical guidelines rather than strict cut-offs. MetaChlam’s accuracy is also inherently dependent on the composition and completeness of its reference databases. The current landscape of *Ct* diversity is shaped by existing sequencing efforts, which are often geographically biased. Limited sequencing data from underrepresented regions may overlook significant genomic variation, particularly for strains like LGV, where sampling bias might have masked within-lineage diversity (Büttner et al. 2025). As sequencing efforts expand, emergence of novel *Ct* strains or greater within-lineage heterogeneity may necessitate periodic MetaChlam database updates to maintain classification accuracy. Without such updates, divergent strains could be misclassified or missed entirely, limiting the pipeline’s ability to capture the full spectrum of *Ct* diversity.

To evaluate MetaChlam beyond controlled datasets in real-world scenarios amidst the complexities of mixed microbial communities, we applied the pipeline to evaluate public metagenomic samples by screening over 100,000 SRA submissions using PebbleScout. MetaChlam analysis of the top 500 datasets revealed that only 158 samples were likely to truly contain *Ct*. Each *Ct*-positive sample had a minimum 5% genome coverage marked by Pebblescout. The majority (150 samples) of these samples originated from well-documented infection sites (or mice studies, 9/150), including the human eye and vaginal microbiomes, consistent with known *Ct* epidemiology (Price et al. 2016). Interestingly, a notable subset of human gut-derived samples was also identified, raising the possibility of rectal infection or transient gut presence of *Ct*, though these findings need further investigation.

A small number of *Ct*-positive samples were detected in unexpected environments: pig guts, an aquarium and deep-sea hydrothermal vents. Given that mis-identification of species in metagenomes can occur through imprecision in typing tools—a famous example being false positive identification of *B. anthracis* and *Y. pestis* in the New York City subway (Afshinnekoo et al. 2015; Petit et al. 2018), we took extra efforts to confirm these findings. In all cases, we found strong evidence that these were real *Ct* reads present, rather than false positive classifications. First, individual reads had high (>99%) sequence identity to the reference *Ct* genomes. Second, putative *Ct* reads were evenly distributed when mapped to a reference genome. Third, assembled contigs were placed within the known *Ct* phylogeny, rather than being outliers, and the *Ct* lineage inferred from the phylogenies also matched the MetaChlam assignment.

While MetaChlam’s ability to achieve high confidence identification of *Ct* reads reduces the likelihood of false positives, the biological context of the samples must always be considered to avoid misinterpretation of results. Detection of organisms outside their expected ecological range in metagenomic datasets can reflect either true but rare biological occurrence or the introduction of contaminant DNA during sample collection, processing, or sequencing. Although it is not entirely implausible that *Ct* could be detected in environments such as the pig gut, given the presence of the closely related species *C. suis*, there is currently no established ecological or experimental evidence supporting *Ct* infection in these settings. In light of the strong genomic evidence supporting the presence of *Ct*-derived reads, but the absence of ecological context supporting true infection, these findings are most consistent with contamination. This highlights an important consideration for metagenomic analyses that even well-characterized human-associated pathogens such as *Ct* can appear in unrelated datasets, and rigorous validation combined with careful interpretation is required to distinguish genuine biological signals from contamination.

The detection of *Ct* in aquarium and undersea volcanic samples is even more difficult to reconcile with known biology. If taken at face value, these observations would imply either an unrecognized host range or an alternative ecological niche for *Ct*, neither of which is currently supported by independent evidence. However, the most likely explanation is that of contamination, either occurring when the material was sampled or during preparation and sequencing in the laboratory. The presence of human reads in each of these samples could be evidence of a contaminating source. However, *Ct* is not a typical laboratory contaminant, as it is usually present at low abundance in clinical samples unless it is grown in the lab, which requires specialized culture conditions for growth and development. The absence of *Ct* in other samples from the same sequencing projects also argues against widespread or systematic contamination. Taken together, these observations suggest that *Ct* detection in these datasets likely reflects isolated contamination events rather than true environmental occurrence. More broadly, these findings highlight that even well-characterized human-associated pathogens can appear in unrelated metagenomic datasets, underscoring the need for rigorous validation and cautious interpretation of unexpected signals.

## Conclusion

This study highlights the use of average nucleotide identity (ANI) as a robust genomic metric for strain-level differentiation. By establishing a 99.75% ANI threshold, we effectively classified *Chlamydia trachomatis* (*Ct*) into its four epidemiologically significant strains—LGV, P-UA, NP-UA, and Ocular. ANI’s precision in resolving phylogenetic relationships aligns with findings from other bacterial species, reinforcing its value as a universal approach for strain classification. Building on this, we developed MetaChlam, a metagenomic pipeline that integrates results from four tools - LINtax, StrainGE, StrainScan, and Sourmash. MetaChlam capitalizes on custom databases to reliably identify *Ct* strains even in complex metagenomic samples. Its application to public datasets uncovered *Ct* presence in unexpected environments, such as pig gut microbiomes and extreme habitats, validated through complementary ANI and phylogenetic analyses. While MetaChlam’s performance is shaped by the current representation of *Ct* diversity in its databases, it offers a powerful framework for strain characterization and pathogen surveillance. As genomic sequencing expands, incorporating new data will further enhance its accuracy and utility, supporting efforts to understand *Ct* evolution, transmission, and public health impacts.

## Methods

### Comparative genomics of *Chlamydia trachomatis* genomes

All publicly available *Ct* genomes were downloaded from the NCBI Assembly database (accessed March 2025) and filtered for quality using the assembly statistics reported by NCBI. Genomes with completeness over 98%, contamination below 2.5%, and N50 value above 50,000 were retained. The filtered dataset with 89 genomes was supplemented with 20 reference genomes representing the various *ompA* sequence types (Olagoke et al. 2025). The resulting set of assembled genomes were used for core-genome analysis using *PIRATE* (version 1.0.5) (Bayliss et al. 2019), following gene annotations performed with *BAKTA* (version 1.9.2) (Schwengers et al. 2021) under default settings. The resulting core gene alignment was used as input for *IQ-TREE2* (version 2.3.0) (Minh et al. 2020) to infer a maximum-likelihood phylogenetic tree using automated model selection and 1,000 ultrafast bootstrap replicates. Tree visualization was performed with the *ggtree* (Yu 2022) package in R.

A distance-based tree was generated by calculating pairwise ANI values with *pyani* (version 0.2.12) (Pritchard et al. 2016). The resulting ANI matrix was visualized as a heatmap using *gplots* (Warnes et al. 2005) package in R. ANI-based clustering was performed with LINflow (version 1.1.0.6) (Tian et al. 2021) using default settings to identify intraspecies clusters. These clusters were named and mapped onto the distance-based tree to establish strain and substrain-level ANI thresholds. A side-by-side comparison of core-genome and distance-based trees was visualized using the *phytools* (Revell 2012) package in R. Additionally, MLST typing was performed using the command-line version of *pyMLST* (version 2.1.6) (Biguenet et al. 2023).

### Simulating metagenomic reads

Simulated metagenomic datasets were generated using *InSilicoSeq* (version 1.6.0) (Gourlé et al. 2019) with the default parameters using the ‘basic’ Illumina error model. For each simulated metagenome, one million paired-end reads, 125 bp in length, were generated by mixing reads derived from *Ct* reference genomes with reads derived from the human reference genome (GRCh38.p14, hg38). *Ct* reads were included at varying proportions to simulate a range of *Ct* abundances and corresponding read depths. These simulated datasets were used to evaluate detection limits, strain-level resolution, and tool performance under controlled conditions.

### MetaChlam: Nextflow pipeline for *Ct st*rain identification from metagenomes

MetaChlam is a reproducible bioinformatic workflow implemented in Nextflow pipeline for the identification and classification of *Ct* from metagenomic Illumina paired-end sequencing datasets. Raw reads serve as input and are processed in parallel by four complementary classifiers:

- StrainGE, which combines k-mer analysis and read alignment to determine the closest matching reference genome, estimate relative abundance, and compute an ANI-based similarity score (ACNI). (van Dijk et al. 2022)
- StrainScan, which uses a tree-based k-mer indexing structure to optimize strain identification accuracy and computational efficiency. It clusters genomes with 95% Jaccard similarity and applies the elastic net model for strain detection and relative abundance prediction. (Liao et al. 2023)
- Sourmash, which utilizes MinHash sketching for rapid sequence comparison. Reference genomes were sketched with k-mer size 31 and a scaling factor of 1000, and sample sketches were compared against the reference database using the “sourmash gather” function. (Titus Brown and Irber 2016)
- LINtax, which adapts Kraken2 (Wood et al. 2019) classification with ANI-informed LIN taxonomy to enable intra-species classification. Customized *Ct*-specific databases were constructed using LINflow-defined clusters (Sharma 2023).

Customized databases tailored to detect and differentiate *Ct* strains were built for all classifiers using the 109 genome set described previously.

The four classifiers operated in parallel and their outputs were aggregated and cross-validated to ensure robustness in the detection and strain typing of *Ct*. MetaChlam reports the detected *Ct* strain(s), the total number of classified *Ct* paired reads, and classifier-level agreement. When sufficient read support was available, classified *Ct* reads were optionally subjected to *ompA* typing using KMA (1.4.15) (Clausen et al. 2018). Final reports for each sample included strain identification, total classified reads, and optionally *ompA* sequence types.

The MetaChlam pipeline is available on Github: https://github.com/parul-sharma/MetaChlam.

### Evaluation using clinical metagenomic data

MetaChlam was evaluated using clinical samples from the NCBI Bioproject PRJNA826539 (Bommana et al. 2022; Joseph et al. 2023). This dataset comprises host-depleted Illumina paired-end metagenomes sequenced from *Ct*-infected human samples. Three samples (362V, 98V, and 192V) were selected for evaluation and downloaded from the Sequence Read Archive (SRA) under accession numbers SRR18765403, SRR18765393, and SRR18765399 respectively. These samples were used to assess MetaChlam’s performance on real-world clinical data and to validate detection thresholds derived from simulated metagenomes.

### Large-scale screening of public metagenomic data

PebbleScout web service (Shiryev and Agarwala 2024) was used to efficiently screen NCBI SRA datasets. *Ct* reference genome (strain D/UW-3/CX, accession NC_000117.1) was used as a query against the metagenome database (queried November 2024). A filtered subset of putative *Ct+* samples were downloaded and processed with the MetaChlam pipeline using the same parameters for all datasets. For the validation of detection of samples from unexpected sources, reads classified as *Ct* were aligned against reference genomes using blastn (version 2.16.0) (Camacho et al. 2009). Classified reads were extracted and assembled into metagenome-assembled genomes (MAGs) using spades (version 4.0.0) (Prjibelski et al. 2020) with the ‘meta’ option. Phylogenetic trees were also constructed for these cases using the 20 *Ct* reference genomes (Olagoke et al. 2025) employing the previously described steps.

## Acknowledgments

We thank Dr Ryan Kelly (University of Washington) and Dr Anna-Louise Reysenbach (Portland State University) for helpful correspondence and clarification regarding the sampling context and interpretation of their published metagenomic datasets.

